# Knowledge, Attitude and Practices towards COVID-19 Guidelines among Students in Bangladesh

**DOI:** 10.1101/2021.03.07.433083

**Authors:** Bezon Kumar, Susmita Dey Pinky, Ashike Md. Nurudden

## Abstract

This paper explores the level of knowledge, attitude and practices of COVID-19 guidelines among the students in Bangladesh. In achieving this objective, this paper uses primary data collected from 1822 students and three different Likert scales and a one-way ANOVA test are used to assess knowledge, attitudes, and practice (KAP) scores and mean differences with respect to different variables. The research reveals that the majority of students have a higher level of knowledge and a positive attitude towards the COVID-19 guidelines. In contrary, only 0.22 percent students show strong compliance towards COVID-19 guidelines while majority students (60.54 percent) have rather a poor adherence which is an alarming finding. Reopening the educational institutions in Bangladesh is, therefore, not advisable yet and reinforcing the preventive measures through campaigns and online discussion to persuade people to follow the preventive guidelines is highly recommended to contain this global disease.

**What We Already Know:**

- Majority papers revealed the KAP of general people or medical related people.
- Majority people have moderate level of knowledge towards COVID-19.
- No research is found on KAP of the students in Bangladesh.

**What This Article Adds:**

- The KAP of COVID-19 guidelines among the students in Bangladesh.
- Majority students have the higher level of knowledge towards COVID-19 guideline among students.
- Only 0.22 percent students show strong compliance towards COVID-19 guidelines which recommends not reopening educational institutions now.

## Introduction

The recent outbreak of respiratory illness caused by a novel (new) coronavirus named (SARS-CoV-2) that was first detected in Wuhan City, Hubei Province, China in December 2019 has kept ravaging the world by spreading to at least 213 countries attributing 12 million deaths and 54 million confirmed cases (worldometer, 2020). As a consequence, the World Health Organization (WHO) declared it a global pandemic on 12 March 2020 due to its high infectious rate with tremendous transmission dynamics (Gumbrecht and Howard, 2020). SARS-CoV-2 is the third coronavirus to emerge in the human population in the past two decades, following the outbreaks of SARS-CoV in 2002 and MERS-CoV in 2012.

Since COVID-19 is an emerging and rapidly evolving situation, even after multiple trials to prove the efficacy of potential drugs that can cure the disease, promising progress seems still far-off (Maragakis, 2020). Hence, following COVID-19 basic infection prevention guidelines as per WHO protocols are the cornerstones of reducing the transmission and they are:

- Practicing social-distance (At least 6 feet away from others)
- Maintaining hand hygiene (Frequent hand washing with soap-water or alcohol-based sanitizer)
- Use of facemasks in public settings.
- Sneezing on elbows
- Quarantining of the exposed individuals to COVID-19 (CDC, 2020).

Ever since the first case was recorded in Bangladesh on March 8, 2020, the exponential rise of case number and death rates have made Bangladesh as one of the worst-hit countries in the world. Up to 23 November, global COVID-19 cases crossed 54 million, wherein Bangladesh reported 0.45 million infected cases and deaths (Coronatracker Bangladesh, 2020).

In light of flattening the coronavirus curve, a major number of schools, colleges and universities all over the world have suspended or cancelled all their face-to-face campus activities and rapidly transitioned to online learning instead. With no exception, the government of Bangladesh has also announced to close all the educational institutions on 16 March 2020 and extended the closure till 19 December 2020 (The Daily Star, November 13, 2020). Meanwhile, Bangladesh has adapted distance learning and many institutions are conducting online classes and tests according to their logistics support and choices. This new norm, however, is creating difficulties for both the students and teachers. On one hand, a great number of students aren’t equipped with adequate devices or a strong broadband network and so, they have to depend on unstable mobile data which are costly as well. On the other hand, many of the teachers lack the required technical skills to conduct online classes. Furthermore, both the teachers and students are yet to be at ease in interacting through screens rather than the traditional face-to-face communication (Tariq and Fami, 2020)

Regarding the public exams, the education ministry has already cancelled the Higher Secondary Certificate (HSC) and its equivalent examinations, which were scheduled to start from 01 April 2020 (TBS, October 7, 2020). Primary school certificate and junior school certificate exams were cancelled earlier too due to Coronavirus situation (Bangla News, 11 August, 2020). Most importantly, the pandemic-induced academies closure has significantly affected the public universities of Bangladesh taking them back in the loops of session-jams which they were fighting against to reverse in the prepandemic period. Most undergraduate students have already lost an academic calendar year, yet there is no sight of reopening the universities in foreseeable future (Anwar, et al., 2020)

After three months of apparent strict containment, the offices, garment factories, courts, shopping malls, public transports all have reopened gradually in order to move the wheel of the economy. Only the educational institutes have remained closed, with no specific decisions taking in an account yet. This, in consequence, rose uncertainty among a group of students compelling them to protest against Government’s decision on extending the closure of the universities in different sites of social media whereas the other group is content with the pace of online classes and support the decision (Shovon, 2020).

It is a well-known fact that to address a public health issue, such as Covid-19 global crisis, people’s knowledge and awareness play a crucial role as adherence to the preventive measures depend on the level of the understanding about the disease and grave consequences are followed by if the preventive controls are not taken properly (Kumar and Pinky, 2020). Again, students of secondary education (grade 6-10), higher secondary and university going students constitute a set of the population whose lives have witnessed a major change throughout this pandemic leaving their academic calendars in disarray. Assessing the knowledge, perspectives and practice toward COVID-19 guidelines among students is, therefore, vital since their perception and practice will have a significant impact on the spread of a pandemic once the educational institutions are open. Similarly, student’s deeper insight into the existing knowledge, responsive attitude and healthy precautionary practice towards the guidelines may curb the infection rate of coronavirus breaking the important chain of transmission in the community (Elmer, et al., 2020).

Thus far, no study has been carried out on assessing the knowledge, awareness and practice towards the COVID-19 guidelines taking students of Bangladesh as the target population. Hence, this study is pivotal as it will denote the adherence to the given protocols by the WHO and as a result, will determine the risks of reopening the educational institutes in near future. On that account, the results of this study will have significant implications for policymakers and further project planners as well.

## Materials and Methods

### Study Design and Sample Selection

This is a cross-sectional study conducted among the students of Secondary (grade 6 to 10), Higher-Secondary (grade 11 to 12) and Tertiary (Undergraduate and Master’s) level in Bangladesh. This paper did not consider primary (grade 1 to 5) students as they are unable to take decisions on their own and generally don’t use the Internet for responding to this survey questionnaire. Amid this pandemic, collecting data from the field through face to face interview was impossible due to social distancing measures, restricted movement and lockdowns. Hence, data were collected online using Google Forms via a self-reported questionnaire and following snowball sampling, the link of the questionnaire was sent to the students of different institutions who forwarded it to others. Moreover, it was shared in social media platforms, namely Facebook, LinkedIn, Twitter, WhatsApp, Messenger so that the sample size becomes larger as the larger the sample size, the higher the external validity and the greater the generalizability of the study (Cavana, et al., 2001). From the latest statistics, it is found that Bangladesh has about 18405709 students from secondary to tertiary level (BANBEIS, 2018). To achieve the study objectives and sufficient statistical power, the representative sample size was calculated with a sample size calculator (RAOSOFT, 2020) and using a margin of error of ±4 percent, a confidence level of 99 percent, a 50 percent response distribution, and 18405709 students, the sample size calculator reached at 1,037 respondents.

Based on the guidelines for the community of COVID-19 by World Health Organization (WHO), the self-reported questionnaire was developed. Initially, the questionnaire was drafted in English and later was translated into Bengali for the online questionnaire. The questionnaire consisted of five primary sections and on the first section, respondents were clearly informed about the background and objectives of the study. They were also apprised that they have full rights to withdraw their response at any point of the survey without giving a reason and that all information and opinions provided would be anonymous and confidential. Respondents who agreed to participate in the study were then instructed to complete the questionnaire. The second section of the questionnaire gathered particulars on socio-demographic features of the respondents while the third section asked for participants’ knowledge of COVID-19 guidelines and the fourth and fifth section collected information on participants’ attitudes, and practices of COVID-19 guidelines, respectively.

The online survey was conducted from 10 October 2020 to 16 November 2020, and finally, 1822 data were collected. The inclusion criteria to participate in the study were being a Bangladeshi student of Secondary to tertiary level, having internet access and voluntary participation. After collection, data were edited, sorted and coded for analysis.

### Measurement of Students’ Knowledge, Attitude and Practices of COVID-19 Guidelines

Respondents were asked ten statements regarding knowledge of COVID-19 guidelines to respond as either true or false or don’t know option. Among ten statements, five statements were given correct and the rest five statements were given wrong. Regarding correct statements, false and don’t know responses were given a score of zero, and true answers were assigned a score of one. In the case of wrong statements, likewise, true and don’t know responses were given a score of zero whereas false answers were assigned a score of one. The total score for knowledge ranged from zero to 10 with scores o to 5 indicating a lower level of knowledge of COVID-19 while scores 6 to 7 and 8 to 10 suggesting a moderate and higher level of knowledge, respectively. Items were evaluated for internal reliability using Cronbach’s *α.* Here, Cronbach’s alpha coefficient was 0.66, implying internal reliability.

In the section on attitudes, scores were calculated based on the respondents’ answers to each attitudinal statement, 1 = undecided, 2 = disagree, and 3 = agree. Scores were calculated by averaging respondents’ answers to the six statements. Total scores ranged from 6 to 18 with scores 6 to 9 denoting lower or negative attitudes while 10 to 14 and 15 to 18 signifying moderate and higher or positive attitudes, respectively. The Likert scales were assessed for internal reliability using Cronbach’s *α.* Herein, Cronbach’s alpha coefficient was 0.64, marking internal reliability.

Regarding practices of COVID-19 guidelines, respondents were asked to respond “always” or “frequently” or “sometimes” or “never” to the items. A score of 3 was given to always while 2, 1 and 0 were given to frequently, sometimes and never. The total score ranged from zero to 36 with scores 0 to 18 indicating a lower level of practices, scores 19 to 27 and 28 to 36 stating a moderate and higher level of practices, respectively. The Likert scales were assessed for internal reliability using Cronbach’s *α.* Cronbach’s alpha coefficient was 0.80, reflecting internal reliability.

### Empirical Methods

This study employed primarily descriptive statistics to tabulate the frequency of social and demographic features of the students. One-way analysis of variance (ANOVA) was used to assess differences in mean values for knowledge, attitudes, and practice (KAP) scores with respect to different factors. The overall mean differences were estimated using a Bartlett test as the scores were continuous (Wetzels, et al., 2012 and O’brien, 1979). All analyses were conducted using SPSS 24 software.

## Results and Discussion

### Social and Demographic Features of the Students

Table 1 shows that among the participants, 53.20 percent of students were male while 44.10 percent were female. While in terms of age groups, the majority of the participants were between (21 to 25 years) and (16 to 20 years), representing 53.95percent and 26.51 percent. The highest qualification of more than 76.70 percent of participants was undergraduate level or above. Regarding residence, more than 51.70 percent of the participants were from urban areas and another 27.70 percent belonged to rural areas. In terms of the duration of using internet per day, more than half of the participants opined between (2 to 3 hours) and (more than 3 hours), representing 20.00 percent and 53.70 percent, respectively. On the other hand, 35.20 percent of the students had the habit of watching TV/reading Newspaper (less than 1 hour) per day, followed by another 16.10 percent who didn’t watch TV/read Newspaper regularly.

**Table 1:**
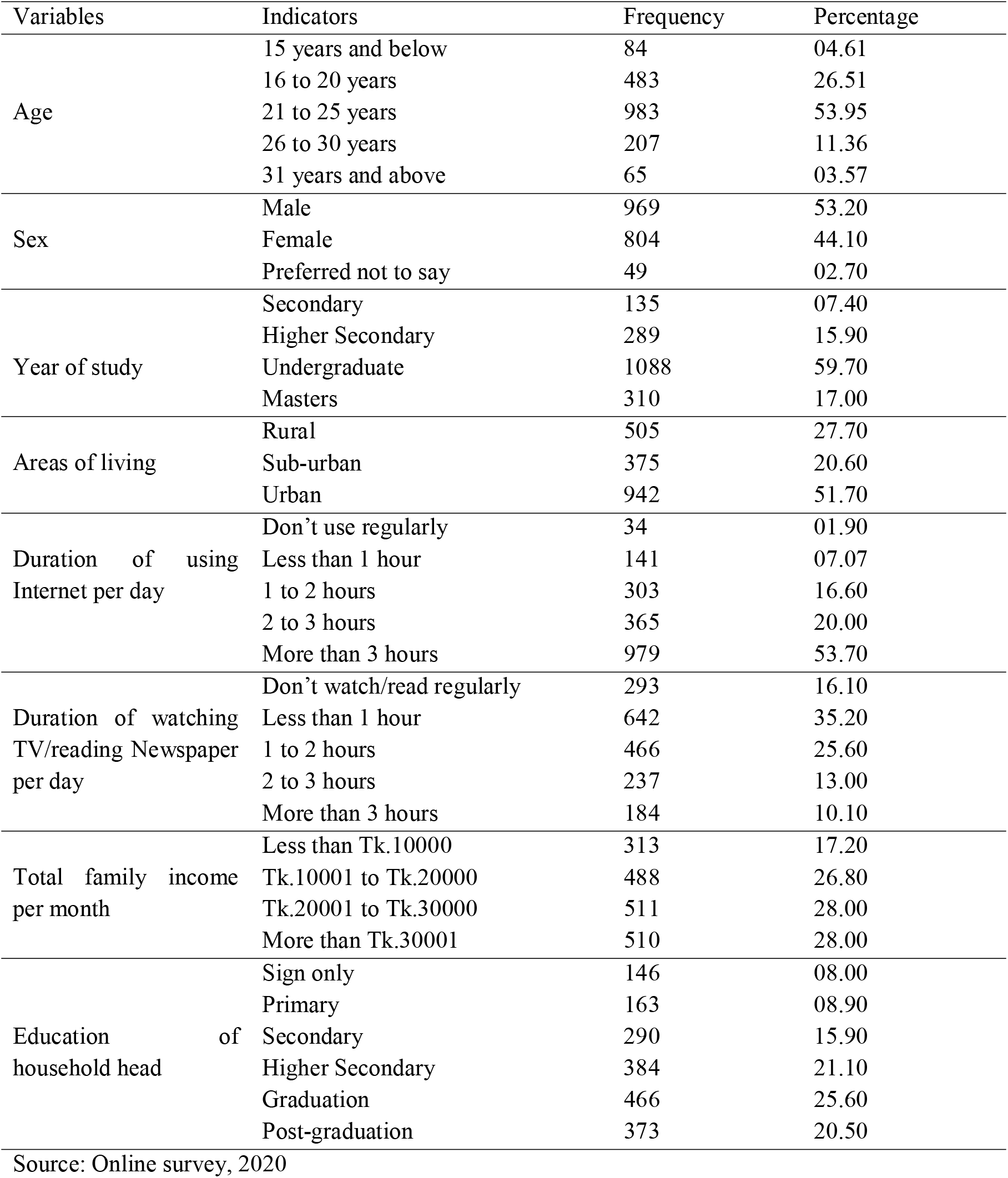
Social and Demographic Features of the Students

**Table 2:** Students’ Responses to Statements regarding Knowledge of COVID-19 Guidelines

| Statements | True | False | Don't Know |
| --- | --- | --- | --- |
| Maintaining 2 feet distance among one another is enough to prevent droplet transmission | 848 (46.5) | 888 (48.7) | 86 (4.7) |
| Wearing a mask is mandatory for the public | 1625 (89.2) | 161 (8.8) | 36 (2.0) |
| Washing hands for 10 seconds with soap-water is enough to kill the virus | 559 (30.7) | 1129 (62.0) | 134 (7.4) |
| Only mouth is the routes of entry of the Coronavirus | 518 (28.4) | 1158 (63.6) | 146 (8.0) |
| Self-isolation is important if someone develops any flu-like symptoms | 1473 (80.8) | 249 (13.7) | 100 (5.5) |
| It is not necessary for children and young adults to take measures to prevent the infection by the COVID-19 virus | 551 (30.2) | 1128 (61.9) | 143 (7.8) |
| Avoiding visits to crowded places can be a way to prevent high rates of droplet transmission | 1587 (87.1) | 148 (8.1) | 87 (4.8) |
| Wearing a surgical mask can fully protect someone from getting coronavirus | 612 (33.6) | 993 (54.5) | 217 (11.9) |
| After being in a public place, after nose-blowing, coughing or sneezing, people must wash their hands with soap and water, or use hand sanitizer containing at least 60percent alcohol, for at least 20 seconds | 1506 (82.7) | 164 (9.0) | 152 (8.3) |
| Pregnant women, elderly people and adults with chronic disease should take extra precautions against coronavirus | 1604 (88.0) | 117 (6.4) | 101 (55.5) |
| Note: Values inside parenthesis shows percentage of total |  |  |  |
| Source: Online survey, 2020 |  |  |  |

### Students’ Responses to Statements regarding Knowledge of COVID-19 Guidelines

A large portion of participants (89.2 percent) answered that wearing a mask is mandatory for the public. The majority of the participants were also aware that avoiding visits to crowded places can be a way to prevent high rates of droplet transmission and self-isolation is important if someone develops any flu-like symptoms, with 87.1 percent and 80.8 percent, respectively. A high portion of the participants (88.0 percent) identified that pregnant women, elderly people and adults with chronic disease are more vulnerable and they should take extra precautions against coronavirus. In addition, about 82.7 percent reported that people must wash their hands with soap and water, or use hand-sanitizer containing at least 60percent alcohol, for at least 20 seconds, after being in a public place, after nose-blowing, coughing or sneezing.

However, nearly 33.6 percent of the participants did not know that wearing a surgical mask can’t fully protect someone from getting affected by coronavirus. Almost 30.2 percent of the students thought that it is not necessary for children and young adults to take measures to prevent the infection by the COVID-19 virus-which is totally wrong. According to WHO, anyone of any age level can be affected by COVID-19. Moreover, about one-quarter of the participants (28.4 percent) regarded mouth as the only route of entry of the Coronavirus whereas respiratory tract, fecal-oral and body fluids all are the possible routes of viral transmission and infection (Li et al. 2020).

Unfortunately, 46.5 percent of the participants affirmed that maintaining 2 feet distance among one another is enough to prevent droplet transmission whereas the appropriate distance should be of six feet. Data from our study showed that 30.7 percent of the participants thought washing hands for only 10 seconds with soap-water is enough to kill the virus when in fact, hands should be washed for at least 20 seconds. The above findings shows that there is some incertitude about issues, such as wearing masks, washing hands and regarding social distancing.

### Students’ Responses to Attitudinal Statements regarding COVID-19 Guidelines

Students’ attitudinal responses are shown in Table 3 where among the participants, around 70 percent showed a positive attitude towards limiting the spread of COVID-19. From the participants’ responses, isolation and self-quarantining for 14 days and maintaining social distancing everywhere ranked first, with 88.9 percent and 86.4 percent of respondents agreeing to these respectively. Wearing mask was ranked second with around 77.8 percent responses of the participants. Unexpectedly, 18.1 percent of the participants didn’t agree with the statement that wearing a mask is effective as a preventive measure. Approximately 74.2 percent of the participants believed that Bangladesh may experience a second wave in winter and the last attitude to limit the spread of COVID-19 identified from participants was the closure of educational institutions with 69.3 percent while 24 percent of the respondents thought it otherwise.

**Table 3:** Students’ Responses to Attitudinal Statements regarding COVID-19 Guidelines

| Statements | Agreed | Not Agreed | Undecided |
| --- | --- | --- | --- |
| Bangladesh is in a good position in controlling COVID-19 | 522 (28.6) | 1123 (61.6) | 177 (9.7) |
| Wearing a mask is effective as a preventive measure | 1417 (77.8) | 329 (18.1) | 76 (4.2) |
| Social distancing is necessary to maintain everywhere | 1574 (86.4) | 178 (9.8) | 70 (3.8) |
| Isolation and self-quarantining for 14 days is important to safeguard others | 1619 (88.9) | 142 (7.8) | 61 (3.3) |
| Bangladesh may experience a second wave in winter | 1352 (74.2) | 224 (12.3) | 246 (13.5) |
| Keeping educational institutions closed can prevent COVID-19 explosion | 1263 (69.3) | 437 (24) | 122 (6.7) |

Note: Values inside parenthesis shows percentage of total
Source: Online survey, 2020

The Bangladesh Government has already taken action in controlling COVID-19 including the lockdown, closure of educational and social organizations, awareness raising activities all over the country and suspension of all domestic and international transportations. In spite of that, majority 61.6 percent of the participants didn’t agree that Bangladesh is in a good position in controlling COVID-19.

### Students’ Responses to Statements regarding Practice of COVID-19 Guidelines

Drawing the data from Table 4, almost 68 percent of participants receded from visiting restaurants/cafes, 50.9 percent avoided from shaking hands and almost 55 percent avoided the crowded place. Unfortunately, the current study showed that still 15.9 percent of students always and 29.5 percent frequently visit crowded place nowadays. In the same way, they visited restaurants/cafes always or frequently, with 7.8 percent and 24.8 percent, respectively. About 78 percent of participants kept themselves updated by following the recent guidelines for COVID-19 regularly. In relation to participants’ practice toward washing hands or foods, 54.4 percent confirmed they always wash hands with soap and water/rubbing alcohol for at least 20 seconds, 70.1 percent confirmed of washing fruits and vegetables well before eating always.

**Table 4:** Students’ Responses to Statements regarding Practice of COVID-19 Guidelines

| Statements | Always | Frequently | Sometimes | Never |
| --- | --- | --- | --- | --- |
| I go out of home amid COVID-19 pandemic | 360 (19.8) | 705 (38.7) | 643 (35.3) | 114 (6.3) |
| I wear a mask when leaving home | 1169 (64.2) | 442 (24.3) | 178 (9.8) | 33 (1.8) |
| I touch the surface of the mask when taking it off | 416 (22.8) | 462 (25.4) | 462 (25.4) | 482 (26.5) |
| I interact with people nowadays | 326 (17.9) | 659 (36.2) | 547 (30.0) | 290 (15.9) |
| I use public transport for commuting | 444 (24.4) | 535 (29.4) | 399 (21.9) | 444 (24.4) |
| I maintain a physical distance of at least 3 feet while being out | 524 (28.8) | 577 (31.7) | 512 (28.1) | 209 (11.5) |
| I visit any crowded place nowadays | 289 (15.9) | 537 (29.5) | 570 (31.3) | 426 (23.4) |
| I avoid some cultural behaviors like shaking hands | 927 (50.9) | 440 (24.1) | 336 (18.4) | 119 (6.5) |
| I wash your hands with soap and water/rubbing alcohol for at least 20 seconds | 992 (54.4) | 486 (26.7) | 281 (15.4) | 63 (3.5) |
| I visit restaurants/cafes meanwhile | 142 (7.8) | 452 (24.8) | 587 (32.2) | 641 (35.2) |
| I keep updates and follow the recent guidelines for COVID-19 | 786 (43.1) | 629 (34.5) | 344 (18.9) | 63 (3.5) |
| I wash fruits and vegetables well before eating | 1277 (70.1) | 319 (17.5) | 188 (10.3) | 38 (2.1) |

Note: Values inside parenthesis shows percentage of total
Source: Online survey, 2020

However, the practice of touching the surface of the mask while taking it off is still controversial. 22.8 percent always do it in a proper way but the rest never do so. Even though around 89 percent of the participants favor wearing a mask while leaving home, 11 percent still did not consider this significant. In addition, 6.5 percent of participants never avoid cultural behaviors like shaking hands, followed by another 18.4 percent who sometimes avoid that. In this study, only 28.8 percent admitted maintaining a physical distance of at least 3 feet while being out, still, 11.5percent never felt that necessity. Likewise, 24.4 percent participants always use public transport for commuting, followed by another 29.4 percent who use frequently. Lastly, a great level of bad practice came into light as majority of the students’ always or frequently went out of home amid COVID-19 pandemic, despite the closure of all educational institutions.

### Level of Students’ Knowledge, Attitude and Practice regarding COVID-19 Guidelines

The level of students’ knowledge, attitude and practice regarding COVID-19 Guidelines is presented in the following table.

Table 5 shows that most of the students 49.78 percent had a higher level of knowledge regarding COVID-19 Guidelines and another 26.67 percent had moderate level of knowledge. On the other hand, 23.55percent had relatively a lower level of knowledge regarding COVID-19 Guidelines. Concerning attitudes, 64.27 percent showed a high level of positive and optimistic attitude towards COVID-19, followed by another 30.57 percent who had a moderate level of positive and optimistic attitude. High-level of knowledge and optimistic attitude among the students, however, was an expected finding as the epidemiological survey was conducted during the later part of the pandemic. Knowledge of the disease is considered the first stepping stone to any health awareness activity since knowing the causes and transmission of a disease accelerates developing positive and vigilant attitude, which invariably results in further good practices. Unfortunately, the findings of the present study contradict with the above notion. In spite of having adequate knowledge and positive attitude, only 0.22 percent had a high level practice, 39.24 percent had a moderate level practice and 60.54 percent had a lower level of practice regarding COVID-19 Guidelines.

**Table 5:**
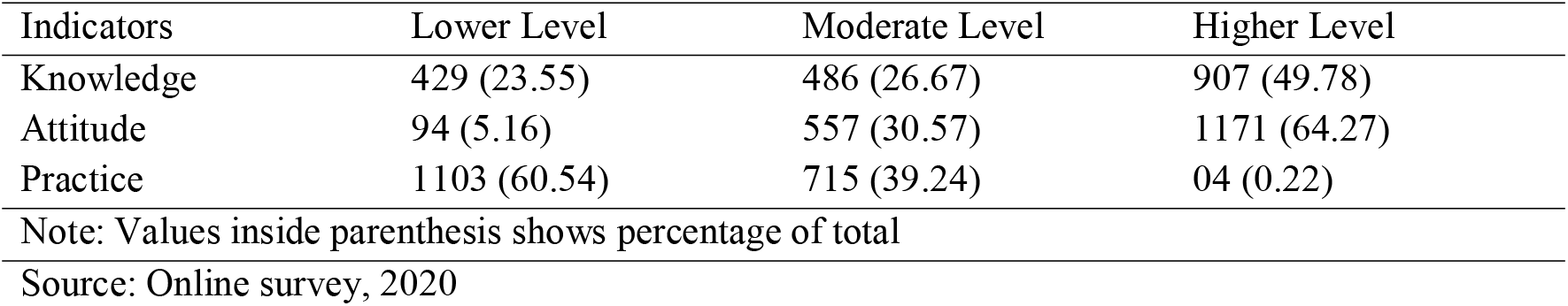
Level of Students’ Knowledge, Attitude and Practice regarding COVID-19 Guidelines

| Indicators | Lower Level | Moderate Level | Higher Level |
| --- | --- | --- | --- |
| Knowledge | 429 (23.55) | 486 (26.67) | 907 (49.78) |
| Attitude | 94 (5.16) | 557 (30.57) | 1171 (64.27) |
| Practice | 1103 (60.54) | 715 (39.24) | 04 (0.22) |

Note: Values inside parenthesis shows percentage of total
Source: Online survey, 2020

### Comparison of Social and Demographic Features, and Mean KAP Score

Table 6 reveals that there is a significant positive correlation between knowledge and attitude, which indicates that a better level of awareness was reflected in their attitude. Interestingly, the correlation between students’ knowledge, attitude and practice differs in a greater margin. Drawing from Table 6, the better practice was found amongst the (16 to 20 years) and (21 to 25 years) age group. Similarly, knowledge and attitude towards COVID-19 guidelines were found higher among them compared to the young or aged group.

**Table 6:** Comparison of Social and Demographic Features, and Mean KAP Score

| Variables | Indicators | Knowledge |  | Attitude |  | Practice |  |
| --- | --- | --- | --- | --- | --- | --- | --- |
|  |  | Mean | F | Mean | F | Mean | F |
| Age | 15 years and below | 6.96 |  | 14.38 |  | 22.74 |  |
|  | 16 to 20 years | 7.07 |  | 15.25 |  | 23.39 |  |
|  | 21 to 25 years | 7.43 | 10.50* | 14.76 | 3.88* | 23.03 | 5.96* |
|  | 26 to 30 years | 6.70 |  | 15.05 |  | 21.34 |  |
|  | 31 years and above | 6.20 |  | 15.08 |  | 20.74 |  |
| Sex | Male | 7.14 |  | 14.71 |  | 21.47 |  |
|  | Female | 7.32 | 10.30* | 15.28 | 23.94* | 24.57 | 58.13* |
|  | Preferred not to say | 7.19 |  | 13.02 |  | 21.49 |  |
| Year of study | Secondary | 6.56 |  | 14.19 |  | 22.28 |  |
|  | Higher Secondary | 6.51 | 19.01* | 15.34 | 6.16* | 21.89 | 3.28** |
|  | Undergraduate | 7.35 |  | 14.87 |  | 23.08 |  |
|  | Masters | 7.50 |  | 15.01 |  | 23.11 |  |
| Areas of living | Rural | 7.29 |  | 15.22 |  | 23.34 |  |
|  | Sub-urban | 6.80 | 7.83* | 14.00 | 29.20* | 21.39 | 12.83* |
|  | Urban | 7.28 |  | 15.12 |  | 23.14 |  |
| Duration of using Internet per day | Don't use regularly | 6.47 |  | 14.26 |  | 19.88 |  |
|  | Less than 1 hour | 6.38 |  | 14.82 |  | 21.83 |  |
|  | 1 to 2 hours | 6.28 | 39.67* | 14.20 | 9.44* | 21.78 | 8.58* |
|  | 2 to 3 hours | 6.91 |  | 14.81 |  | 22.45 |  |
|  | More than 3 hours | 7.71 |  | 15.21 |  | 23.55 |  |
| Duration of watching TV/reading Newspaper per day | Don't watch/read regularly | 7.51 |  | 15.26 |  | 22.47 |  |
|  | Less than 1 hour | 7.46 |  | 15.08 |  | 23.11 |  |
|  | 1 to 2 hours | 7.04 | 10.92* | 14.68 | 5.91* | 23.25 | 3.66* |
|  | 2 to 3 hours | 6.67 |  | 14.35 |  | 21.54 |  |
|  | More than 3 hours | 6.74 |  | 15.14 |  | 23.07 |  |
| Total family income per month | Less than Tk.10000 | 7.17 |  | 14.89 |  | 23.50 |  |
|  | Tk.10001 to Tk.20000 | 7.01 |  | 14.59 |  | 22.69 |  |
|  | Tk.20001 to Tk.30000 | 7.02 | 6.28* | 14.94 | 4.68* | 21.94 | 6.46* |
|  | More than Tk.30001 | 7.51 |  | 15.22 |  | 23.46 |  |
| Education of household head | Sign only | 6.93 |  | 14.58 |  | 21.65 |  |
|  | Primary | 7.12 |  | 14.22 |  | 21.76 |  |
|  | Secondary | 7.04 |  | 14.77 |  | 21.72 |  |
|  | Higher Secondary | 7.12 | 8.01* | 14.62 | 7.59* | 22.77 | 7.90* |
|  | Graduation | 6.95 |  | 15.22 |  | 23.22 |  |
|  | Post-graduation | 7.78 |  | 15.39 |  | 24.21 |  |

Note: \*, \*\* indicate 1 percent, and 5 level of significance, respectively
Source: Online survey, 2020

Female showed slightly better knowledge, attitude and practice towards national guidelines than males. This can be explained by different gender-related activities in social and in the family roles. Undergraduate students and who had master’s degrees were much aware of COVID-19 guidelines. The analysis reveals that students’ knowledge, attitude and practice may be increased if their level of education increased and it is significant at 5 percent level of significance. The rational explanation may be that the higher the educational level, the more their depth and extent of knowledge and the more positive attitude towards the guidelines as well. The study finds that rural people had more knowledge, optimistic attitude and good practices towards COVID-19. This may be because they were most likely to belong at home, there’s no such public gathering or activities in rural areas.

Concerning sources of knowledge on COVID-19, it is not surprising that internet and online media took the lead, though traditionally TV/newspaper remains the leading media of awareness in this country. It is noteworthy that the use of the internet and online media is increasing among the students constantly. With regard to the duration of using internet per day, participants’ knowledge, attitude and practice increased with the duration of their internet use. This indicates that a significant portion of participants in the survey was largely influenced by media information. Due to limited access to internet or online resources, a number of respondents have poor knowledge, negative attitudes and inappropriate preventive practices towards COVID-19.

The study finds no particular relation between students’ habit of reading newspaper/watching TV and their knowledge, attitude and practice. The analysis showed that students’ with higher income had better knowledge and attitude; nevertheless, students who had lower family income had a good practice towards COVID-19 guidelines. The rational explanation may be that students having the higher family income have the higher opportunities to spend on educational purposes but then they also have more chances to visit restaurants/tourist spots so they were less likely to follow COVID-19 guidelines. However, students’ who belong to an educated family had greater knowledge, attitude and practice towards COVID-19 guidelines. This is significant at 1 percent level of significance and can be interpreted by the fact that educated parents are much aware of health and diseases.

Finally, the study finds that it’s the practice, not the knowledge/attitude; where the emphasis should be given. The findings here may be helpful to inform policymakers and health professionals, on further public health interventions, policies and significant decision making.

### Conclusion and Policy Recommendations

This paper principally investigates two research questions. First, what is the level of students’ knowledge, attitude and practices of COVID-19 guidelines in Bangladesh? Second, what factors are associated with the level of students’ knowledge, attitude and practices of COVID-19 guidelines? In finding out the solutions to these questions, this paper used primary data collected from 1822 students and several empirical methods and pinpointed some significant findings.

Firstly, this research ascertains that the majority of students (49.78 percent) had the higher level of knowledge of COVID-19 guidelines while 26.67 percent and 23.55 percent students had a moderate and lower level of knowledge, respectively. Regarding attitude, a majority of students (64.27 percent) held a very positive perspective towards the COVID-19 guidelines, followed by another (30.57 percent) who shows moderate optimism towards it. Surprisingly yet, only 0.22 percent students had a higher level of practices of COVID-19 guidelines whereas majority students (60.54 percent) mentioned the lower level of practices. Secondly, this paper also reveals that the higher the students’ study year, Internet use, family income, and education of household head, the higher the extent of students’ knowledge, attitude and practices of COVID-19 guidelines.

Based on this paper’s findings, it is recommended to keep the educational institutions closed in Bangladesh since almost no students adequately practice the COVID-19 guidelines and so, if the institutions reopen now, these schools, colleges might become reservoirs of virus in the probable upcoming second wave of the pandemic in winter. Since the time and budget of this paper were constraints, this paper did not consider a larger sample size or employ more relevant empirical methods. Thus, the findings of this paper may not represent the actual scenario of Bangladesh and therefore, it recommends researchers to carry out a further in-depth study on this issue excluding these constraints.

## Acknowledgement

Authors of this paper are thankful to BK School of Research for providing technical assistance. Authors are also thankful to the Research Assistants for their valuable assistance, without which this research would not be possible.

## Author Contributions

BK conceived and designed the study. Both BK and SDP made the questionnaire in English and translated into Bengali and set in Google Forms. While SDP wrote introduction, BK wrote methodology and analyzed data, and AMN wrote results and discussions. BK reviewed and edited the whole paper which was finally approved by all authors as the final version.

## Data Availability

All data analyzed during this study are available online as a supplementary file.

